# Effects of predictive and incentive value manipulation on sign- and goal-tracking behavior

**DOI:** 10.1101/767095

**Authors:** Cristina E. Maria-Rios, Christopher J. Fitzpatrick, Jonathan D. Morrow

## Abstract

When a neutral stimulus is repeatedly paired with an appetitive reward, two different types of conditioned approach responses may develop: a sign-tracking response directed toward the neutral cue, or a goal-tracking response directed toward the location of impending reward delivery. Sign-tracking responses have been postulated to result from attribution of incentive value to conditioned cues, while goal-tracking reflects the assignment of only predictive value to the cue. We therefore hypothesized that sign-tracking would be more sensitive to manipulations of incentive value, while goal-tracking would be more responsive to changes in the predictive value of the cue. We tested sign- and goal-tracking before and after devaluation of a food reward using lithium chloride, and tested whether either response could be learned under negative contingency conditions that precluded any serendipitous reinforcement of the behavior that might support instrumental learning. We also tested the effects on sign- and goal-tracking of blocking the predictive value of a cue using simultaneous presentation of a pre-conditioned cue. We found that sign-tracking was sensitive to outcome devaluation, while goal-tracking was not. We also confirmed that both responses are Pavlovian because they can be learned under negative contingency conditions. Goal-tracking was completely blocked by a pre-conditioned cue, while sign-tracking was only partially reduced. These results indicate that sign- and goal-tracking follow different rules of reinforcement learning and suggest a need to revise current models of associative learning to account for these differences.

## Introduction

Modern theories of associative learning are a crowning achievement of experimental psychology. These theories were developed as a way to comprehensibly and systematically describe how behavioral responses to sensory inputs change with experience. Many behavioral patterns have been identified and explained by learning theories, including classical conditioning, extinction, conditioned inhibition, and blocking. Learning theory has been successfully applied clinically through the development and refinement of effective psychiatric treatments such as cognitive-behavioral therapy, exposure therapy, and contingency management. However, one potential limitation of current theories is that they generally assume all subjects are learning the same information under a given set of experimental conditions, and therefore the same set of rules can be used to explain learning and behavior for all individuals. Of course, even in a simple Pavlovian learning experiment, not all subjects will respond in exactly the same way. In many cases, individual variation manifests as differences in the magnitude of the response to stimuli, for example freezing to a greater or lesser degree to a stimulus associated with footshock (Bush, Sotres□Bayon, & LeDoux, 2007). This type of variation is often simply dismissed as “noise”, though some have argued that characterizing these behavioral differences could be important for understanding “abnormal” psychiatric pathology (Galatzer-Levy, Bonanno, Bush, & LeDoux, 2013). In at least some cases, individual differences in conditioned behaviors go beyond just the magnitude of the responses. More qualitative differences in the type of behavioral responses displayed could suggest a more fundamental difference between individuals in the type of information being learned.

For example, during a Pavlovian conditioned approach (PCA) procedure in which a neutral cue such as a retractable lever (conditioned stimulus; CS), is repeatedly paired with delivery of a food-pellet reward (unconditioned stimulus; US) to a different location, two conditioned responses may develop. One is a “sign-tracking” response in which the subject approaches and interacts with the cue itself, and another is a “goal-tracking” response in which the subject moves away from the cue and towards the site of impeding food delivery whenever the cue is presented (Flagel, Akil, & Robinson, 2009; Tomie & Morrow, 2018). If this procedure is performed on a large number of outbred rats, some individuals will engage in both sign- and goal-tracking, but a large proportion will almost exclusively learn to perform only one or the other conditioned response (Meyer et al., 2012).

There is some evidence that sign- and goal-trackers differ not only in the form of their behavioral response to the CS, but also in the type of information they are gleaning from the association between the CS and US. For example, after PCA training sign-trackers are much more likely to learn a new operant response and work for the presentation of the CS than goal-trackers when tested on a conditioned reinforcement procedure (T. E. Robinson & Flagel, 2009; Villaruel & Chaudhri, 2016). The way in which sign-trackers interact with the CS can also differ depending on the nature of the US; for example eating behaviors will be directed toward a CS paired with solid food, but drinking behaviors will be directed toward a CS paired with liquid (Jenkins & Moore, 1973). This indicates that for sign-trackers, but not goal-trackers, the CS may take on some of the motivational properties of the US and thereby gains its own incentive value. Interestingly, goal-tracking behaviors are quicker to extinguish than sign-tracking when the US is consistently omitted (Ahrens, Singer, Fitzpatrick, Morrow, & Robinson, 2016; Beckmann & Chow, 2015; Fitzpatrick, Geary, Creeden, & Morrow, 2019). Because extinction is based on removing the predictive relationship between the CS and US, the resistance of sign-tracking to extinction might suggest that predictive value does not influence sign-tracking behavior the same way that it influences goal-tracking. These findings at least raise the possibility that different individuals could be following different rules of associative learning, even though they are being tested under identical experimental conditions.

Based on evidence outlined above, we hypothesize that goal-tracking behavior is mainly influenced by the predictive value of the CS, but sign-tracking is primarily influenced by incentive value that is ultimately derived from the US. We therefore predict that goal-tracking will be more sensitive to manipulations involving only the predictive value of the CS, whereas sign-tracking will be more sensitive to manipulations of incentive value. As a way of isolating these two properties in order to test this hypothesis, we used a Pavlovian blocking procedure to selectively manipulate the predictive value of the CS, and a devaluation procedure to change its incentive value.

Blocking is a counterintuitive Pavlovian learning phenomenon that, at least in part, the Rescorla-Wagner model of associative learning was developed to explain. In this learning paradigm, originally described by Leo Kamin in the late 1960s, the appearance of a CS that has already been associated with the US will prevent, or “block”, learning to a second CS that is presented at the same time (Kamin, 1967a, 1967b). The Rescorla-Wagner model accounts for this by saying that the previously trained CS eliminates the prediction error “surprise” signal that is necessary for any further associative learning to occur in response to the US (Rescorla & Wagner, 1972). The US retains its full incentive value in this scenario, but no new conditioned response develops because the presence of a pre-trained stimulus deprives the new CS of its predictive value.

In contrast, devaluation procedures remove the incentive value of the US while leaving the predictive relationship between CS and US intact. Devaluation of a food US is commonly achieved by pairing the food with a compound such as lithium chloride (LiCl), which will induce nausea and abdominal discomfort that mimics food poisoning (Holland & Rescorla, 1975; Holland & Straub, 1979). Many species, including rats, have evolved to be very sensitive to this type of learning and will quickly lose all interest in consuming food that has been paired with these aversive interoceptive effects. Thus, if our hypothesis is correct, we would expect sign-tracking responses to be more resistant to a Pavlovian blocking procedure than goal-tracking, whereas goal-tracking should be more resistant than sign-tracking to devaluation with LiCl. Because Pavlovian responses are thought to be more sensitive to devaluation than operant responses (Balleine & O’Doherty, 2010), we also performed a negative contingency experiment to confirm that both sign- and goal-tracking are Pavlovian responses, as opposed to “superstitious” operant responses that can sometimes occur due to serendipitous pairings of a behavior with a US (Skinner, 1948).

## Methods

### Animals

Adult male Sprague Dawley rats (275-300g) were purchased from Harlan Laboratories and Charles River Laboratories in order to maximize diversity of behavioral phenotypes (Fitzpatrick et al., 2013). All experimental groups were counterbalanced to ensure equal numbers of subjects from each vendor. Rats were maintained on a 12:12-hr light/dark cycle, and food and water were available ad libitum for the duration of experimentation. Rats were acclimatized to the housing colony for two days prior to handling. All procedures were approved by the University Committee on the Use and Care of Animals (University of Michigan; Ann Arbor, MI).

### Drugs

Lithium chloride (LiCl; Acros Organics / Thermo Fisher Scientific, Inc.; Waltham, MA) was used. LiCl was dissolved in 0.9% sterile saline to make a 0.3 M solution (pH = 7.38-7.42), and 0.9% sterile saline was used as a vehicle control.

### Behavioral Testing Apparatus

Sixteen modular operant conditioning chambers (24.1 cm width × 20.5 cm depth x 29.2 cm height; MED Associates, Inc.; St. Albans, VT) were used for Pavlovian conditioning. Each chamber was located in a sound-attenuating cubicle equipped with a ventilation fan to provide ambient background noise. For testing of PCA, each chamber was equipped with a food magazine, a retractable lever (counterbalanced on the left or right side of the magazine), and a red house light on the wall opposite of the magazine. For experiment 3, a tone generator was also located above the magazine, and a second lever was present such that for this experiment a lever was located both to the left and right of the magazine. The magazine contained an infrared sensor to detect magazine entries, and the levers were calibrated to detect lever deflections in response to 10 g of applied weight. Whenever a lever was extended into the chamber, an LED mounted inside the lever mechanism illuminated the slot through which the lever protruded. Number, latency, and probability of lever presses and magazine entries were recorded automatically (ABET II Software; Lafayette Instrument; Lafayette, IN).

### Experiment 1: Pavlovian Conditioned Approach and Devaluation Procedures

For two days prior to the start of training rats were familiarized with banana-flavored pellets (45 mg; Bio-Serv; Frenchtown, NJ) in their home cages. Rats were then placed into the test chambers for one pretraining session during which the red house-light remained on but the lever was retracted. Twenty-five food pellets were delivered on a variable time (VT) 30-s schedule (i.e., one pellet was delivered on average every 30 s, but actually varied 0-60 s). Rats were not food deprived during at any point during experimentation. Next, rats underwent seven daily sessions of PCA training. Each trial during a PCA training session consisted of presentation of the illuminated lever (conditioned stimulus; CS) into the chamber for 8 s on a VT 90-s schedule (i.e., time randomly varied 30-150 s between CS presentations). Retraction of the lever was immediately followed by the response-independent delivery of one pellet (unconditioned stimulus; US) into the magazine. The beginning of the next inter-trial interval commenced immediately after pellet delivery. Each test session consisted of 25 trials of a CS-US pairing. All rats consumed all the pellets that were delivered.

Following PCA training, rats were placed in conditioning chambers for four 10-min devaluation sessions during which with the red house light was on and 25 banana-flavored pellets were already present in the pellet magazine. Rats could freely feed during the duration of the four devaluation sessions. Immediately following the end of the session, rats were administered LiCl (0.3 M) at a volume of 5 mL/kg (i.p.). At the end of each session, the number of pellets consumed was recorded for each rat. Twenty-four hours after the last devaluation session, rats underwent a devaluation test session during which sign- and goal-tracking conditioned responses were measured. This test session was identical to a PCA training session except it was under extinction conditions (i.e., rats received 25 lever-CS presentations in the absence of the food pellet delivery).

### Experiment 2: Pavlovian conditioned approach procedure with a negative contingency

Pretraining was performed as in Experiment 1. Next, rats underwent twelve daily sessions of PCA training with a negative contingency. Each trial during these PCA training sessions consisted of presentation of the illuminated lever (CS) into the chamber for 8 s on a VT 90-s schedule. As in Experiment 1, training sessions consisted of 25 CS presentations. However, in this experiment there was a negative contingency such that retraction of the lever was only followed by food pellet delivery (US) into the magazine if no conditioned response was detected. In other words, lever presses or magazine entries during the CS-period cancelled delivery of the food pellet at the offset of the CS-period. Number, latency, and probability of lever presses and magazine entries were recorded automatically (ABET II Software; Lafayette Instrument). In addition, ceiling-mounted cameras were used to record behavior during training sessions and measure approach to the lever or the magazine during each CS-period. All rats consumed all the pellets that were delivered, and pellets delivered during a training session were recorded for each rat, though due to the negative contingency not every rat was administered 25 pellets during a session.

### Experiment 3: Pavlovian blocking

Pretraining was performed as in Experiments 1 and 2. Rats were divided into two counterbalanced groups before the study began: control (CTL) or blocking (BLK). For the BLK group, Phase 1 of training (pre-blocking) consisted of six days of PCA training as described in Experiment 1 with presentation of one lever (L1; counterbalanced left and right of pellet magazine) preceding food pellet delivery. For the CTL group, the same protocol was performed using an 8-s tone (4 KHz, 80dB) as the CS instead of a lever. For both CTL and BLK groups, Phase 2 (blocking) consisted of six more days of PCA training, but during this phase each US delivery was preceded by the simultaneous extension of two levers (L1 and L2) into the chamber for 8 s. Twenty-four hours after the last session of Phase 2, rats from both CTL and BLK groups underwent one post-blocking test in which the L2 lever, which had previously only been encountered during Phase 2 of training, was extended into the chamber. No food pellets were delivered during the post-blocking test.

## Statistical Analysis

In Experiment 1, PCA behavior was scored using an index that incorporates the number, latency, and probability of lever presses (sign-tracking conditioned response) and magazine entries (goal-tracking conditioned response) during CS presentations within a session (Meyer et al. 2012). Briefly, we averaged the response bias (i.e., number of lever presses and magazine entries for a session; [lever presses – magazine entries] / [lever presses + magazine entries]), latency score (i.e., average latency to perform a lever press or magazine entry during a session; [magazine entry latency – lever press latency]/8), and probability difference (i.e., proportion of lever presses or magazine entries; lever press probability – magazine entry probability). The index score ranges from +1.0 (absolute sign-tracking) to −1.0 (absolute goal-tracking), with 0 representing no bias. The average PCA index scores of Session 6-7 were used to classify rats as STs (score ≥ 0.5), GTs (score ≤ −0.5), and intermediate responders (−0.5 < score < 0.5). For the BLK group in Experiment 3, this index score was determined during the Phase 1 sessions 5 and 6 based on the conditioned responses to L1. Because the CTL group in Experiment 3 had a tone during Phase 1 that did not allow measures for sign-tracking behavior, their index score was determined during Phase 2 based on their sign-tracking behavior to both L1 and L2. In Experiment 2, a modified approach index, similar to the probability difference used in the PCA index, was used to phenotype rats. The proportion of approaches to the lever or magazine in Session 12 was used to generate an index that ranges from +1.0 to −1.0, and rats were classified as STs (score ≥ 0.5) GTs (score ≤ −0.5), and intermediate responders (−0.5 < score < 0.5). Because these studies were designed to detect reductions in conditioned behavior, intermediate responders were excluded from analysis so that the relatively low levels of sign- and goal-tracking from these individuals would not interfere with our ability to detect behavioral effects from the experimental manipulations.

SPSS (Version 24; IBM, Inc.) was used for all statistical analysis. For all linear mixed models, the covariance structure was selected based upon Akaike’s information criterion (i.e., the lowest number criterion represents the highest quality statistical model using a given covariance structure). In Experiment 1, PCA behavior across training sessions were analyzed using a linear mixed model with an autoregressive (AR1) covariance structure with Response (Lever Press and Magazine Entry Number) and Session as factors. For devaluation sessions, pellet consumption was analyzed using a linear mixed model (AR1) with Drug (Saline and LiCl) and Session as factors. For the devaluation test session, behavior was analyzed using a linear mixed model (AR1) with Response, Session (Pre- and Post-Devaluation), and Devaluation (Saline and LiCl) as factors. In addition, behavior was analyzed using difference scores and a two-way analysis of variance (ANOVA) with Response and Drug as factors. Difference scores were calculated by subtracting the number of lever presses or magazines entries from the first trial of the devaluation test from the first trial of Session 7.

In Experiment 2, approach index scores were analyzed using an ANOVA with Phenotype (ST and GT) as a factor. Goal-tracking behavior across the 12 training sessions was analyzed using a linear mixed model (AR1) with Period (CS and non-CS) and Session as factors. Sign-tracking behavior was analyzed using a two-way ANOVA with Response (Approach and Contact) and Session (Session 1 and 12) as factors. Pellet consumption was analyzed using a linear mixed model (AR1) with Phenotype and Session as factors. When appropriate for any statistical test, multiple comparisons were performed using Fisher’s Least Significant Difference (LSD) post hoc test.

For the Phase 1 sessions of Experiment 3, both BLK and CTL PCA behavior across training was analyzed using a linear mixed model with an autoregressive (AR1) covariance structure. For the CTL group their magazine entry number, latency, and probability were analyzed with Session (1-6) as a factor. For the BLK group, lever press and magazine entry number, latency, and probability as well as PCA index score were analyzed with Phenotype (GTs and STs) and Session (1-6) as factors. Same analysis was performed for the CTL group during Phase 2. For STs in the CTL and BLK groups, preferential contact with L1 versus L2 during the last session of Phase 2 was analyzed using paired T-tests. Magazine entry number for GTs were compared between CTL and BLK groups using an independent T-test. For the test sessions, lever press number of STs from both CTL and BLK groups and magazine entry number of GTs from both CTL and BLK in the first trial of the session, were compared to zero using one-sample T-tests. For the BLK group, lever press and magazine entry number from STs and GTs respectively, were compared to those in session six of Phase 1 using paired-sample T-tests. Multiple comparisons were performed using Fisher’s Least Significant Difference (LSD) post hoc test.

## Results

### Experiment 1: Devaluation

Rats underwent seven days of PCA training and were classified as STs, GTs, and intermediate responders based upon the average PCA index scores during Session 6-7. However, only STs (n = 11) and GTs (n = 19) were used for further experimental testing. By definition, STs and GTs differed in their lever press number (effect of Phenotype; F_(1,37.63)_ = 38.04, p = 3.47 × 10^−7^), latency (effect of Phenotype; F_(1,43.69)_ = 84.55, p = 8.89 × 10^−12^), and probability (effect of Phenotype; F_(1,44.41)_ = 117.08, p = 4.93 × 10^−14^) as well as their magazine entry number (effect of Phenotype; F_(1,40.26)_ = 49.09, p = 1.78 × 10^−8^), latency (effect of Phenotype; F_(1,42.86)_ = 79.45, p = 2.58 × 10^−11^), and probability (effect of Phenotype; F_(1,41.67)_ = 54.62, p = 4.23 × 10^−9^). Following the seven daily PCA training sessions, rats underwent four sessions of devaluation training. Rats were divided into the following groups: ST/LiCL (n = 10), ST/Saline (n = 9), GT/LiCl (n = 6), and GT/Saline (n = 5). Treatment with LiCl (n = 16) compared to saline (n = 14) devalued consumption of the banana-flavored pellets (Figure 1A; effect of Drug; F_(1,76.57)_ = 737.1, p = 4.82 × 10^−41^; interaction of Drug x Session; F_(1,138.37)_ = 165.31, p = 1.49 × 10^−45^). Evidence of devaluation was detected in the difference score between conditioned responding on the first CS presentation of the devaluation test subtracted from the seventh (baseline) PCA training session (Figure 1B; interaction of Drug x Phenotype; F_(1,26)_ = 21.88, p = 0.037). Post hoc comparisons revealed that only STs (p = 0.002), and not GTs (p = 0.89), devalued their conditioned responding following exposure to LiCl.

**Figure 1.**
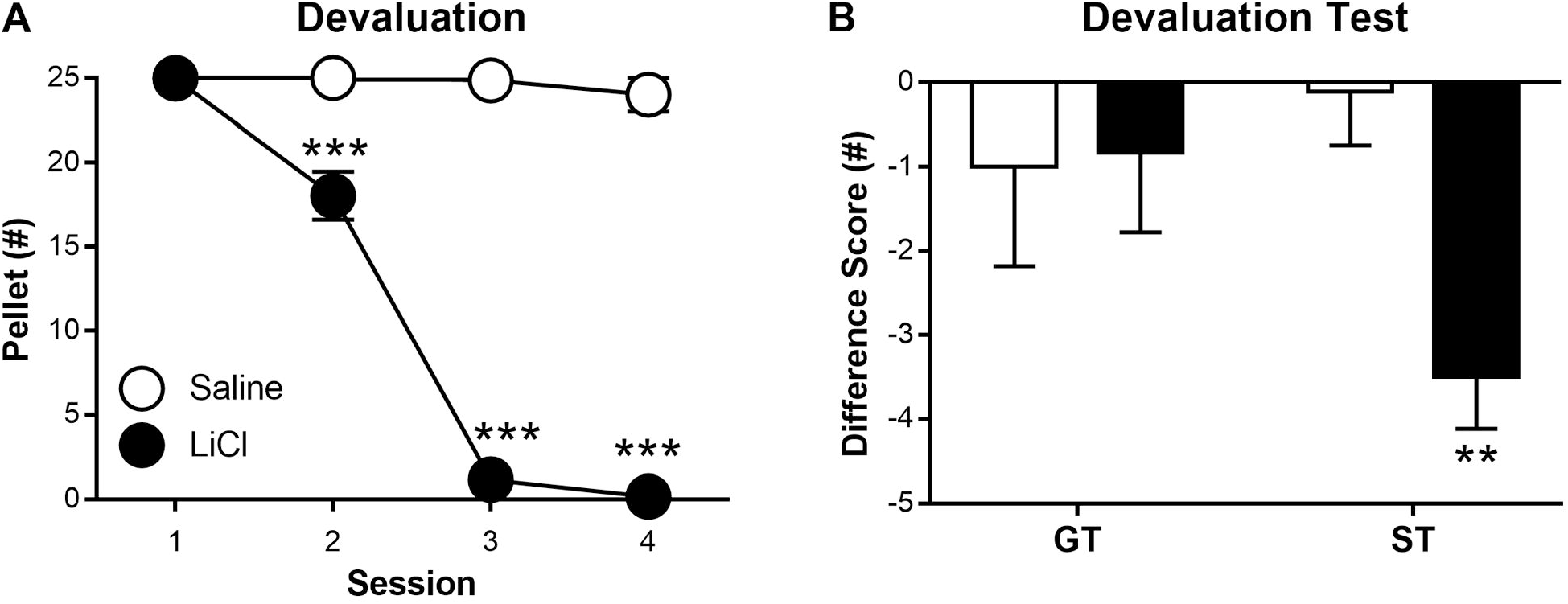
Devaluation. (A) Pellet consumption during four sessions of devaluation training during which rats could freely feed on 25 pellets for 10 min within the conditioning chambers. Immediately following removal from the chambers, rats were injected with 0.3 M LiCl (5 mL/kg, i.p.; n = 25) or saline vehicle (5 ml/kg, i.p.; n = 23). (B) A difference score was calculated for the number of magazine entries in goal-trackers (GTs) and the number of lever presses in sign-trackers (STs) during the first trial of the last PCA training session (Session 7), subtracted from the number during the first trial of the devaluation test. Data are presented as mean ± SEM. ** = p < 0.01, *** = p < 0.001.

### Experiment 2: Negative Contingency

Rats were classified as GTs or STs using their approach index score on Session 12 (Figure 2A; effect of Phenotype; t_20_ = −24.11, p = 2.97 × 10^−16^). During training, STs (n = 10) learned to avoid actual contact with the lever approach but still increased their approach to the lever between Sessions 1 and 12 (Figure 2B; F_(1,60)_ = 25.54, p = 4.34 × 10^−6^). In contrast, GTs (n = 11) learned to magazine-enter during the CS-period compared to the NCS-period, despite the fact that conditioned responding cancelled food pellet delivery (Figure 2C; interaction of Period x Session; F_(11, 81.86)_ = 2.12, p = 0.028). As a result, STs earned more pellets than GTs during the 12 PCA training sessions with a negative contingency (Figure 2D; F_(11, 165.21)_ = 1.93, p = 0.039). Post hoc comparisons revealed that STs earned more pellets than GTs on Sessions 8-12 (p < 0.05).

**Figure 2.**
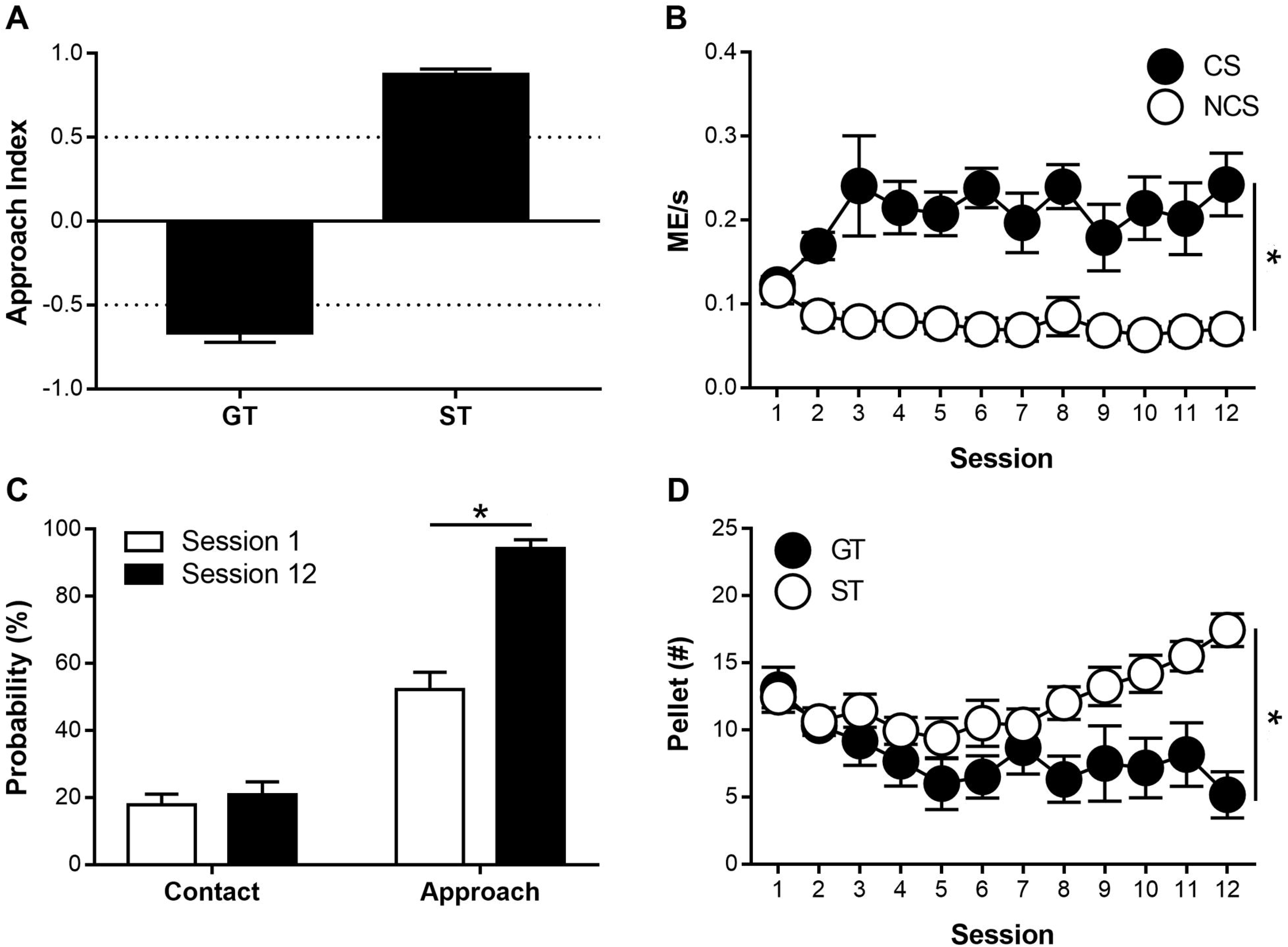
Negative contingency (i.e., lever presses or magazine entries during the conditioned stimulus period cancelled food pellet delivery). (A) Approach index scores from Session 12 that were used to classify rats as sign-trackers (STs) or goal-trackers (GTs). The formula for this score was (# of approaches to the lever - # of approaches to the magazine) / total # of approaches. (B) Magazine entries per second during the conditioned stimulus (CS) periods and non-CS periods, graphed across all 12 training sessions. (C) Probability of sign-tracking behavior (both contacts with and approaches to the lever) on session 1 and session 12 of training. (D) Number of pellets earned by STs and GTs graphed across all 12 training sessions. Data are presented as mean ± SEM. * = p < 0.05.

### Experiment 3: Pavlovian Blocking

Experiment 3 was conducted in 3 phases as diagramed in Figure 3.

**Figure 3.**
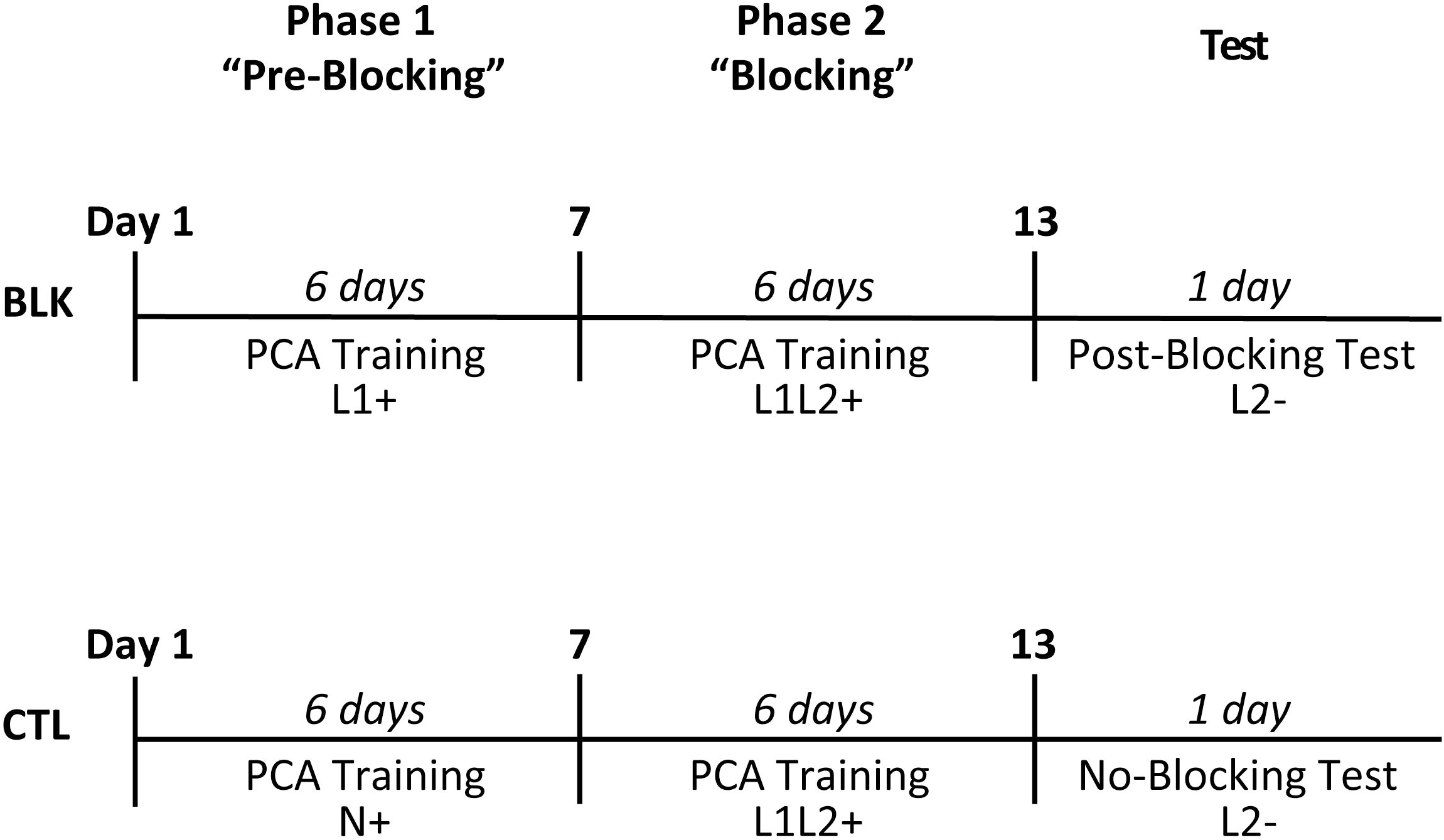
Pavlovian blocking experimental design. During Phase 1 “Pre-Blocking”, rats in the BLK group underwent 6 days of PCA training with a lever (L1+) as the food-reinforced CS, while for rats in the CTL group an auditory tone (N+) served as the reinforced CS during the 6 days of PCA training. During Phase 2 “Blocking”, rats from both the CTL and BLK group underwent 6 days of PCA training with L1 (pre-trained lever for the BLK group) and simultaneous presentation of a new lever L2 forming the combined reinforced CS (L1L2+). Finally, all rats underwent one test session in which L2 was presented with no food reward (L2-). This test session was identified as “Post-Blocking” for the BLK group and “No-Blocking” for the CTL group.

#### Phase 1: Pre-Blocking Sessions

Rats from the BLK group underwent six days of PCA training with a lever CS (L1) and were classified as either STs or GTs according to their PCA index score (intermediate responders were excluded from further analysis). By the end of Session 6, STs (n = 23) and GTs (n = 8), exhibited significantly different lever press number (effect of Phenotype; F_(1,44.833)_ = 88.991, p = 3.246 × 10^−12^), latency (effect of Phenotype; F_(1,40.946)_ = 142.001, p = 6.8489 × 10^−15^), and probability (effect of Phenotype; F_(1,39.000)_ = 294.192, p = 9.1748 × 10^−20^) as well as magazine entry number (effect of Phenotype; F_(1,34.100)_ = 83.114, p = 1.1471 × 10^−10^), latency (effect of Phenotype; F_(1,35.606)_ = 123.392, p = 4.0303 × 10^−13^), and probability (effect of Phenotype; F_(1,40.296)_ = 162.284, p = 1.0273 × 10^−15^). STs and GTs also differed in their PCA index score across the six training sessions (effect of Phenotype; F_(1,34.801)_ = 362.842, p = 5.4812 × 10^−20^). Rats from the CTL group (n = 48) underwent six days of PCA training with an auditory tone CS (N). During training, the rats developed a conditioned response across sessions as evidenced by an increase in magazine entry number (effect of Session; F_(5,209.473)_ = 17.645, p = 1.4094 × 10^−14^), probability (effect of Session; F_(5,213.182)_ = 24.326, p = 2.3945 × 10^−19^) and decrease in latency (effect of Session; F_(5,219.162)_ = 28.117, p = 5.6938 × 10^−22^).

#### Phase 2: Blocking Sessions

After Phase 1, rats from both BLK and CTL groups underwent six days of PCA training with the pre-trained lever (L1) and a simultaneously presented new lever (L2) as the CS. Rats from the CTL group were classified as STs or GTs according to their PCA index score during this Phase 2 (intermediate responders were excluded from further analysis). For classification purposes responses on both L1 and L2 were counted as lever presses. By the end of Session 6, STs (n = 17) and GTs (n = 6) differed significantly in lever press number (effect of Phenotype; F_(1,24.559)_ = 23.686, p = 0.000055), latency (effect of Phenotype; F_(1,26.056)_ = 33.958, p = 0.000004), probability (effect of Phenotype; F_(1,25.625)_ = 76.712, p = 3.5134 × 10^−9^) and magazine entry number (effect of Phenotype; F_(1,29.140)_ = 41.204, p = 4.9691 × 10^−7^), latency (effect of Phenotype; F_(1,30.398)_ = 48.923, p = 8.373 × 10^−8^), and probability (effect of Phenotype; F_(1,31.963)_ = 61.580, p = 5.9888 × 10^−9^). In addition, STs and GTs differed in their PCA index score across the six training sessions (effect of Phenotype; F_(1,27.239)_ = 114.718, p = 2.8735 × 10^−11^). By the last blocking session (Session 6), the number of L1 lever presses was higher than L2 for rats in the BLK group (Figure 4A; paired T-test; t_(22)_ = 3.831076, p = 0.000910) but this preference was not exhibited by STs in the CTL group (Figure 4A; paired T-test; t_(16)_ = 1.100780, p = 0.287278). GTs in the CTL and BLK groups did not differ in their magazine entry number during the last blocking session (Figure 4B; independent T-test; t_(8.801)_ = 1.816064, p = 0.103493).

**Figure 4.**
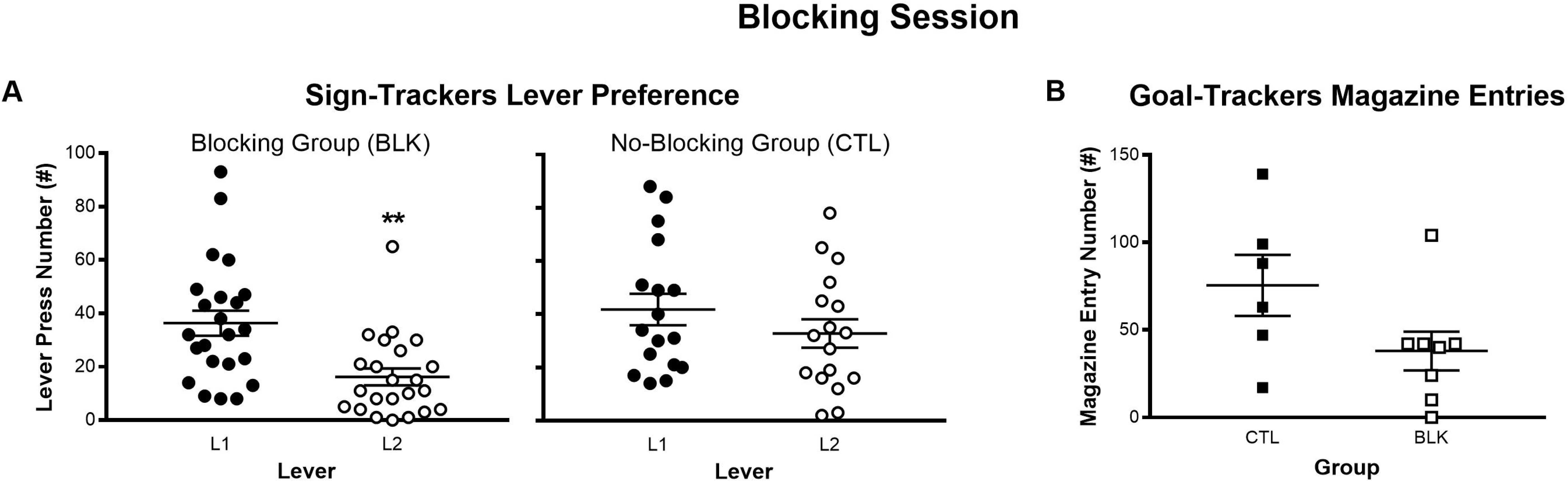
Blocking session. (A) Number of lever presses for levers L1 and L2 during Phase 2 “Blocking”. For the “Blocking” group (BLK) the L1 lever had been previously reinforced in Phase 1, but for the “No-Blocking” control group (CTL) neither lever had been previously encountered. (B) Number of magazine entries during the Phase 2 CS presentations for the BLK and CTL groups. Data are presented as mean ± S.E.M. ** = p < 0.001.

#### Test: Post-Blocking Session

Rats from both the BLK and CTL group underwent one test session in which L2 was presented with no paired food reward. For both STs (n = 23) and GTs (n = 8) in the BLK group, we compared the conditioned responses to Lever 2 (lever presses and magazine entries respectively) on the first trial of this test session to zero (zero was considered as a full blocking effect) as well as to their conditioned responses to Lever 1 during the first trial of the final session of Phase 1 (Pre-Blocking). For STs, lever press number was decreased in this Post-Blocking test session compared to the Pre-Blocking trial (Figure 5A; paired-sample T-test; t_(22)_ = 3.411, p = 0.002507), however lever press number during this test session was significantly different from zero (Figure 5A; one-sample T-test; t_(22)_ = 4.299, p = 0.000291), indicating an incomplete blocking of the sign-tracking response. For GTs, the Post-Blocking magazine entry number was decreased compared to the Pre-Blocking trial (Figure 5B; paired-sample T-test; t_(7)_ = 2.662, p = 0.032394), and magazine entry number during the Post-Blocking test session was not significantly different from zero (Figure 5B; one-sample T-test; t_(7)_ = 1.871, p = 0.103552), indicating complete blocking of the goal-tracking response. In the No-Blocking group that had been previously conditioned with an auditory tone, conditioned responses to Lever 2 during the test session were significantly different from zero for STs’ lever press number (Figure 5C; one-sample T-test; t_(16)_ = 5.215, p = 0.000085) as well as for GTs’ magazine entries (Figure 5C; one-sample T-test; t_(5)_ = 2.712, p = 0.042194), indicating that prior associative learning experiences in general were not sufficient to block either sign- or goal-tracking.

**Figure 5.**
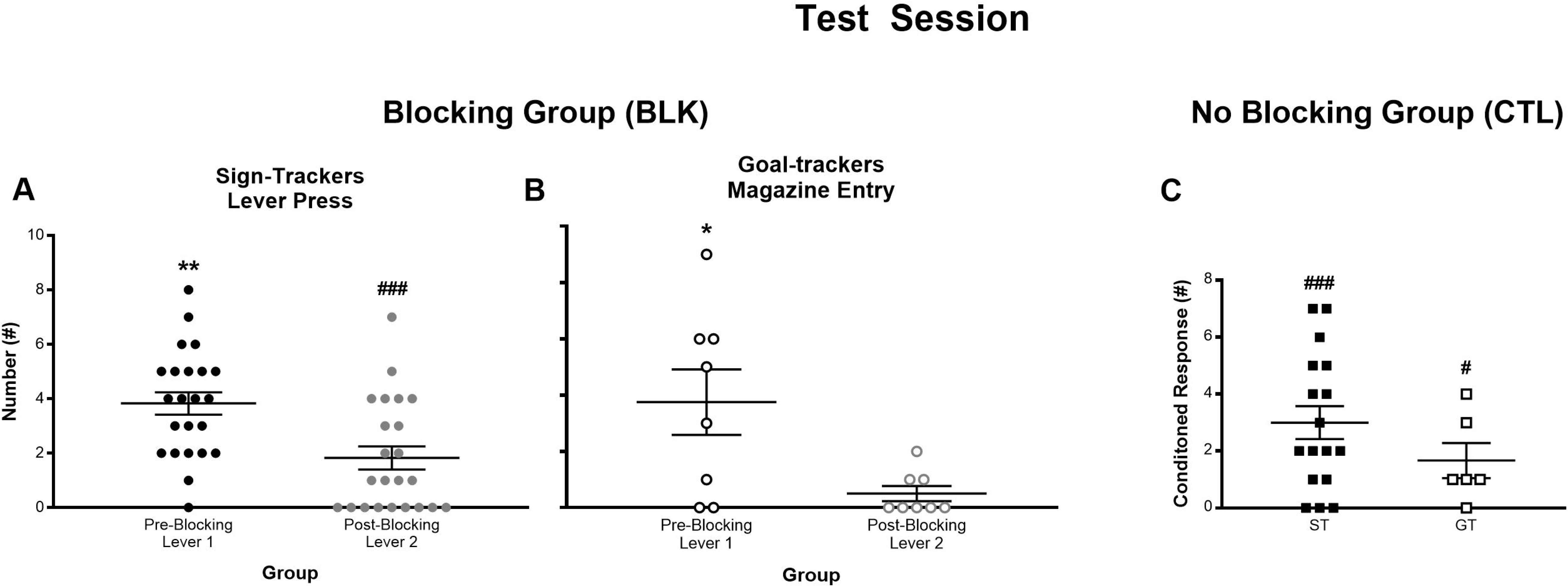
Pre-vs Post-Blocking test session. (A) Number of lever presses for STs in the first trial of the last Pre-Blocking session, and in the Post-Blocking test trial for the “Blocking” (BLK) group. (B) Number of magazine entries during lever presentation for GTs in the first trial of the last Pre-Blocking session, and in the Post-Blocking test trial for the “Blocking” (BLK) group. (C) Number of conditioned responses (lever presses for STs and magazine entries for GTs) during presentation of the L2 lever in the “No-Blocking” test trial for the “No-Blocking” (CTL) control group. Data are presented as mean ± S.E.M. For Pre-versus Post-Blocking paired-sample T-tests: * = p < 0.05, ** = p < 0.01. For one-sample T-tests to detect a significant difference from 0, # = p <0.05, ### = p < 0.0001.

## Discussion

In Experiment 1, we found that sign-tracking is sensitive to outcome devaluation while goal-tracking is not. After subjecting STs and GTs to four sessions of outcome devaluation using lithium chloride, STs significantly reduce their lever press responses in a test session while GTs’ magazine entries remain unaltered during the presentation of the conditioned cue. Experiment 2 confirms that both sign- and goal-tracking are Pavlovian responses rather than “superstitious” instrumental behaviors since both responses develop despite a negative contingency that cancels the delivery of the reward if either response is performed. Interestingly, we found that goal-trackers consistently poke their noses into the food cup even though this behavior results in no reward, while sign-trackers learn to approach the lever without engaging in direct contact with it. In Experiment 3, we found that simultaneous presentation of a previously-trained lever cue was enough to completely block the acquisition of a goal-tracking response to a new lever, but only partially blocked sign-tracking responses to the new lever.

The results of Experiment 1 are in agreement with several older studies that have demonstrated sensitivity of sign-tracking responses to outcome devaluation procedures (Cleland & Davey, 1982; Davey & Cleland, 1984; Holland & Straub, 1979; Holloway & Domjan, 1993). However, until relatively recently no studies have used devaluation procedures to directly compare sign- and goal-tracking on a task that can easily distinguish the two behaviors. Nasser et al. found that individual rats who preferentially sign-track are less likely than goal-trackers to show sensitivity to outcome devaluation on a Pavlovian task, but the effects of devaluation on sign- and goal-tracking behavior were not specifically examined in that study (Nasser, Chen, Fiscella, & Calu, 2015). In more direct contrast to our results, Morrison et al. reported that goal-tracking responses were more sensitive than sign-tracking to outcome devaluation (Morrison, Bamkole, & Nicola, 2015). Though the reason for this discrepancy is unclear, it is possible that subtle methodological differences may have influenced the results. Though the lever CS used by Morrison et al. was essentially the same as that employed in the present study, their US was a liquid sucrose reward delivered along with a tone that had previously been paired with sucrose delivery. Auditory cues paired with food delivery are known to elicit magazine approaches even in sign-trackers (Meyer, Cogan, & Robinson, 2014). Though conditioned responses elicited by tone-CSs are topographically indistinguishable from goal-tracking, they might actually reflect a mixture of the learning elements involved in both sign- and goal-tracking. In fact, very few of the subjects in the Morrison et al. study exhibited behaviors that would qualify them as sign-trackers under the criteria used in our study (Meyer et al., 2012), raising the possibility that at least some of the apparent goal-tracking responses of those subjects actually reflected learning processes that normally underlie sign-tracking behavior. More in keeping with the present study, Derman et al. showed clearly that sign-tracking responses are sensitive to devaluation, though a dearth of goal-trackers in those experiments made it difficult to assess the effects on goal-tracking behavior (Derman, Schneider, Juarez, & Delamater, 2018). On balance, the literature seems to support the conclusion that sign-tracking is readily responds to outcome devaluation, and the current results add to a smaller pool of evidence suggesting that goal-tracking is relatively insensitive to such changes in incentive value.

The differential sensitivity of sign- and goal-tracking to LiCl manipulations is surprising because according to most modern interpretations, Pavlovian responses should always be sensitive to outcome devaluation (Balleine & O’Doherty, 2010). It has been shown that instrumental responses start out as “goal-directed” and are sensitive to devaluation early in training, but with extended training they become “habitual” and insensitive to outcome devaluation (Adams & Dickinson, 1981). This is thought to be because the stimulus is initially associated with the outcome (S-O), and upon stimulus presentation the subject performs a response in order to obtain the desired outcome. Upon more extensive training, the stimulus becomes more directly associated with the response (S-R), and stimulus presentation will then trigger a conditioned response regardless of the incentive value of the outcome. In contrast, Pavlovian learning involves only an association between the conditioned and unconditioned stimuli (S-S^*^), so no such transition to S-R should be possible, and the conditioned response should always remain sensitive to outcome devaluation (Holland, 2008). The apparent resistance of goal-tracking to devaluation could make sense if goal-tracking were not a true Pavlovian response. Even though in the present experiment rats did not need to perform any response in order for food to be delivered, some rats might have entered the food magazine by chance prior to the US and formed a false “superstitious” believe that goal-tracking behavior is required for US delivery. For those animals, goal-tracking would actually be an instrumental response that at some point in training would become insensitive to devaluation. The negative contingency in Experiment 2 controlled for such a possibility because the US was canceled by the goal-tracking response, so the two could never become accidentally associated. The fact that goal-tracking responses developed even under these circumstances indicates that goal-tracking is Pavlovian after all, although it might not follow all the canonical rules of Pavlovian learning (Harris, Andrew, & Kwok, 2013; Holland, 1979). Like previous studies, our results also confirm that sign-tracking is Pavlovian because it is not prevented by a negative contingency, though it does shift to an approach response rather than actual contact with the lever CS (Chang & Smith, 2016; Locurto, Terrace, & Gibbon, 1976; Williams & Williams, 1969). Goal-tracking may have been less amenable to such a shift in strategy because it was recorded by crossing an infrared beam that the rats could not detect, presumably making it difficult for them to discern exactly how close they could approach without canceling the US.

The results from Experiment 3 are largely in concordance with previous reports showing that sign-tracking responses resist blocking from pre-trained levers or auditory cues, while levers readily block goal-tracking to auditory cues (Derman et al., 2018; Holland et al., 2014). It is somewhat difficult to interpret studies that use an auditory tones CS to block a moving lever CS or vice versa because levers are considerably more salient than tones, and thus can command attention and overshadow any predictive information carried by a tone CS. Experiment 3 in the present study avoids this problem of differential attentional salience by using identical levers for blocking. Because we were able to detect a significant amount of both sign- and goal-tracking behavior elicited by the levers, we were able to assess the effects of blocking on both behaviors when trained under identical circumstances. Using identical levers raises additional concerns about generalization of learning due to a reduced ability to discriminate between the levers, but the clear preference of sign-trackers for the pre-trained lever and the evidence of blocking itself in goal-trackers shows that the rats could discriminate between the two stimuli.

The evidence presented here and elsewhere in the literature suggest that sign-tracking is heavily influenced by the incentive value of the US (Jenkins & Moore, 1973; M. J. F. Robinson & Berridge, 2013; T. E. Robinson & Flagel, 2009), while goal-tracking is more sensitive to the predictive relationship between the CS and US (Ahrens et al., 2016; Fitzpatrick et al., 2019). This is not to say that there is a complete dichotomy between the two behaviors; both incentive and predictive value are likely necessary for the acquisition of sign- and goal-tracking. For example, a sign-tracking response will not develop to a cue that is not explicitly paired with reward (T. E. Robinson & Flagel, 2009; Villaruel & Chaudhri, 2016). However, the apparent differential sensitivity of sign- and goal-tracking to manipulations of incentive and predictive values suggests that they reflect two different learning processes that are both normally involved in the acquisition of a Pavlovian conditioned response. Which process comes to dominate the behavioral response likely depends on multiple factors, including those inherent to the individual such as genetic differences (Fitzpatrick et al., 2013; Flagel et al., 2010; Kearns, Gomez-Serrano, Weiss, & Riley, 2006) and early life experiences (Anderson, Bush, & Spear, 2013; Beckmann & Bardo, 2012; Lomanowska et al., 2011), as well as variations in the specific details of the learning task such as physical characteristics of the CS, relative timing and location of the CS and US, etc. (Burns & Domjan, 2001; Fitzpatrick & Morrow, 2016; Holland, 1980; Meyer et al., 2014; Silva, Silva, & Pear, 1992). For example, decreases in the probability of reinforcement tend to increase sign-tracking responses at the expense of goal-tracking (Anselme, Robinson, & Berridge, 2013; Boakes, 1977). Understanding the rules governing this type of learning can have important implications for the prevention and treatment of maladaptive behaviors like pathological gambling and internet gaming disorders, which could be characterized as resulting from reward learning under conditions that favor sign-tracking (Blaszczynski & Nower, 2002; Hellberg, Russell, & Robinson, 2019; Starcke, Antons, Trotzke, & Brand, 2018). Our results could also help to explain some of the difficulty researchers have encountered with reproducing important phenomena such as Kamin’s original blocking effect (Maes et al., 2016). For example, if the cohort being tested happens to be genetically predisposed toward sign-tracking, their responses may prove resistant to blocking, and a cohort predisposed toward goal-tracking may not show much of a devaluation effect. Further attention to these behavioral differences should facilitate future attempts at refining our understanding of associative learning processes.

## Acknowledgements

This work was funded by the University of Michigan Department of Psychiatry (U032826 [JDM]), the Department of Defense National Defense Science and Engineering Graduate Fellowship (CJF), the University of Michigan Predoctoral Fellowship (CJF), the National Science Foundation Graduate Research Fellowship Program (CMR), and the National Institute on Drug Abuse (NIDA; K08 DA037912 [JDM]; R01 DA044960 [JDM]; T32 DA007281 [CJF]).

## References

Adams, C. D., & Dickinson, A. (1981). Instrumental responding following reinforcer devaluation. The Quarterly Journal of Experimental Psychology Section B, 33(2b), 109–121.

Ahrens, A. M., Singer, B. F., Fitzpatrick, C. J., Morrow, J. D., & Robinson, T. E. (2016). Rats that sign-track are resistant to Pavlovian but not instrumental extinction. Behavioural Brain Research, 296, 418–430. https://doi.org/10.1016/j.bbr.2015.07.055

Anderson, R. I., Bush, P. C., & Spear, L. P. (2013). Environmental manipulations alter age differences in attribution of incentive salience to reward-paired cues. Behavioural Brain Research, 257, 83–89. https://doi.org/10.1016/j.bbr.2013.09.021

Anselme, P., Robinson, M. J. F., & Berridge, K. C. (2013). Reward uncertainty enhances incentive salience attribution as sign-tracking. Behavioural Brain Research, 238, 53–61. https://doi.org/10.1016/j.bbr.2012.10.006

Balleine, B. W., & O’Doherty, J. P. (2010). Human and rodent homologies in action control: Corticostriatal determinants of goal-directed and habitual action. Neuropsychopharmacology, 35(1), 48–69. https://doi.org/10.1038/npp.2009.131

Beckmann, J. S., & Bardo, M. T. (2012). Environmental enrichment reduces attribution of incentive salience to a food-associated stimulus. Behavioural Brain Research, 226(1), 331–334. https://doi.org/10.1016/j.bbr.2011.09.021

Beckmann, J. S., & Chow, J. J. (2015). Isolating the incentive salience of reward-associated stimuli: Value, choice, and persistence. Learning & Memory, 22(2), 116–127. https://doi.org/10.1101/lm.037382.114

Blaszczynski, A., & Nower, L. (2002). A pathways model of problem and pathological gambling. Addiction, 97(5), 487–499. https://doi.org/10.1046/j.1360-0443.2002.00015.x

Boakes, R. A. (1977). Performance on learning to associate a stimulus with positive reinforcement. In H. Davis & H. M. B. Hurwitz (Eds.), Operant-Pavlovian interactions (Vol. 67, p. 97). Hillsdale, N.J.□: New York: L. Erlbaum Associates□; distributed by the Halsted Press Division of J. Wiley.

Burns, M., & Domjan, M. (2001). Topography of spatially directed conditioned responding: Effects of context and trial duration. Journal of Experimental Psychology: Animal Behavior Processes, 27(3), 269–278. https://doi.org/10.1037/0097-7403.27.3.269

Bush, D. E. A., Sotres‐Bayon, F., & LeDoux, J. E. (2007). Individual differences in fear: Isolating fear reactivity and fear recovery phenotypes. Journal of Traumatic Stress, 20(4), 413–422. https://doi.org/10.1002/jts.20261

Chang, S. E., & Smith, K. S. (2016). An omission procedure reorganizes the microstructure of sign-tracking while preserving incentive salience. Learning & Memory (Cold Spring Harbor, N.Y.), 23(4), 151–155. https://doi.org/10.1101/lm.041574.115

Cleland, G. G., & Davey, G. C. L. (1982). The effects of satiation and reinforcer develuation on signal-centered behavior in the rat. Learning and Motivation, 13(3), 343–360. https://doi.org/10.1016/0023-9690(82)90014-5

Davey, G. C. L., & Cleland, G. G. (1984). Food anticipation and lever-directed activities in rats. Learning and Motivation, 15(1), 12–36. https://doi.org/10.1016/0023-9690(84)90014-6

Derman, R. C., Schneider, K., Juarez, S., & Delamater, A. R. (2018). Sign-tracking is an expectancy-mediated behavior that relies on prediction error mechanisms. Learning & Memory, 25(10), 550–563. https://doi.org/10.1101/lm.047365.118

Fitzpatrick, C. J., Geary, T., Creeden, J. F., & Morrow, J. D. (2019). Sign-tracking behavior is difficult to extinguish and resistant to multiple cognitive enhancers. Neurobiology of Learning and Memory, 163, 107045. https://doi.org/10.1016/j.nlm.2019.107045

Fitzpatrick, C. J., Gopalakrishnan, S., Cogan, E. S., Yager, L. M., Meyer, P. J., Lovic, V., … Morrow, J. D. (2013). Variation in the form of Pavlovian conditioned approach behavior among outbred male Sprague-Dwley rats from different vendors and colonies: Sign-tracking vs. Goal-tracking. PLOS ONE, 8(10), e75042. https://doi.org/10.1371/journal.pone.0075042

Fitzpatrick, C. J., & Morrow, J. D. (2016). Pavlovian conditioned approach training in rats. JoVE (Journal of Visualized Experiments), (108), e53580. https://doi.org/10.3791/53580

Flagel, S. B., Robinson, T. E., Clark, J. J., Clinton, S. M., Watson, S. J., Seeman, P., … Akil, H. (2010). An animal model of genetic vulnerability to behavioral disinhibition and responsiveness to reward-related cues: Implications for addiction. Neuropsychopharmacology, 35(2), 388–400. https://doi.org/10.1038/npp.2009.142

Galatzer-Levy, I. R., Bonanno, G. A., Bush, D. E., & LeDoux, J. (2013). Heterogeneity in threat extinction learning: Substantive and methodological considerations for identifying individual difference in response to stress. Frontiers in Behavioral Neuroscience, 7. https://doi.org/10.3389/fnbeh.2013.00055

Harris, J. A., Andrew, B. J., & Kwok, D. W. S. (2013). Magazine approach during a signal for food depends on Pavlovian, not instrumental, conditioning. Journal of Experimental Psychology: Animal Behavior Processes, 39(2), 107–116. https://doi.org/10.1037/a0031315

Hellberg, S. N., Russell, T. I., & Robinson, M. J. F. (2019). Cued for risk: Evidence for an incentive sensitization framework to explain the interplay between stress and anxiety, substance abuse, and reward uncertainty in disordered gambling behavior. Cognitive, Affective, & Behavioral Neuroscience, 19(3), 737–758. https://doi.org/10.3758/s13415-018-00662-3

Holland, P. C. (1979). Differential effects of omission contingencies on various components of Pavlovian appetitive conditioned responding in rats. Journal of Experimental Psychology: Animal Behavior Processes, 5(2), 178–193. https://doi.org/10.1037/0097-7403.5.2.178

Holland, P. C. (1980). Influence of visual conditioned stimulus characteristics on the form of Pavlovian appetitive conditioned responding in rats. Journal of Experimental Psychology: Animal Behavior Processes, 6(1), 81–97. https://doi.org/10.1037/0097-7403.6.1.81

Holland, P. C. (2008). Cognitive versus stimulus-response theories of learning. Learning & Behavior, 36(3), 227–241. https://doi.org/10.3758/LB.36.3.227

Holland, P. C., Asem, J. S. A., Galvin, C. P., Keeney, C. H., Hsu, M., Miller, A., & Zhou, V. (2014). Blocking in autoshaped lever-pressing procedures with rats. Learning & Behavior, 42(1), 1–21. https://doi.org/10.3758/s13420-013-0120-z

Holland, P. C., & Rescorla, R. A. (1975). The effect of two ways of devaluing the unconditioned stimulus after first- and second-order appetitive conditioning. Journal of Experimental Psychology: Animal Behavior Processes, 1(4), 355–363. https://doi.org/10.1037/0097-7403.1.4.355

Holland, P. C., & Straub, J. J. (1979). Differential effects of two ways of devaluing the unconditioned stimulus after Pavlovian appetitive conditioning. Journal of Experimental Psychology: Animal Behavior Processes, 5(1), 65–78. https://doi.org/10.1037/0097-7403.5.1.65

Holloway, K. S., & Domjan, M. (1993). Sexual approach conditioning: Tests of unconditioned stimulus devaluation using hormone manipulations. Journal of Experimental Psychology: Animal Behavior Processes, 19(1), 47–55. https://doi.org/10.1037/0097-7403.19.1.47

Jenkins, H. M., & Moore, B. R. (1973). The form of the auto-shaped response with food or water reinforcers. Journal of the Experimental Analysis of Behavior, 20(2), 163–181. https://doi.org/10.1901/jeab.1973.20-163

Kamin, L. J. (1967a). Attention-like processes in classical conditioning (Technical Report No. 5). Hamilton, Ontario: McMaster University.

Kamin, L. J. (1967b). Predictability, surprise, attention, and conditioning (Technical Report No. 13). Hamilton, Ontario: McMaster University.

Kearns, D. N., Gomez-Serrano, M. A., Weiss, S. J., & Riley, A. L. (2006). A comparison of Lewis and Fischer rat strains on autoshaping (sign-tracking), discrimination reversal learning and negative automaintenance. Behavioural Brain Research, 169(2), 193–200. https://doi.org/10.1016/j.bbr.2006.01.005

Locurto, C., Terrace, H. S., & Gibbon, J. (1976). Autoshaping, random control, and omission training in the rat. Journal of the Experimental Analysis of Behavior, 26(3), 451–462. https://doi.org/10.1901/jeab.1976.26-451

Lomanowska, A. M., Lovic, V., Rankine, M. J., Mooney, S. J., Robinson, T. E., & Kraemer, G. W. (2011). Inadequate early social experience increases the incentive salience of reward-related cues in adulthood. Behavioural Brain Research, 220(1), 91–99. https://doi.org/10.1016/j.bbr.2011.01.033

Maes, E., Boddez, Y., Alfei, J. M., Krypotos, A.-M., D’Hooge, R., De Houwer, J., & Beckers, T. (2016). The elusive nature of the blocking effect: 15 failures to replicate. Journal of Experimental Psychology: General, 145(9), e49–e71. https://doi.org/10.1037/xge0000200

Meyer, P. J., Cogan, E. S., & Robinson, T. E. (2014). The form of a conditioned stimulus can influence the degree to which it acquires incentive motivational properties. PLOS ONE, 9(6), e98163. https://doi.org/10.1371/journal.pone.0098163

Meyer, P. J., Lovic, V., Saunders, B. T., Yager, L. M., Flagel, S. B., Morrow, J. D., & Robinson, T. E. (2012). Quantifying Individual Variation in the Propensity to Attribute Incentive Salience to Reward Cues. PLOS ONE, 7(6), e38987. https://doi.org/10.1371/journal.pone.0038987

Morrison, S. E., Bamkole, M. A., & Nicola, S. M. (2015). Sign tracking, but not goal tracking, is resistant to outcome devaluation. Frontiers in Neuroscience, 9. https://doi.org/10.3389/fnins.2015.00468

Nasser, H. M., Chen, Y.-W., Fiscella, K., & Calu, D. J. (2015). Individual variability in behavioral flexibility predicts sign-tracking tendency. Frontiers in Behavioral Neuroscience, 9. https://doi.org/10.3389/fnbeh.2015.00289

Robinson, M. J. F., & Berridge, K. C. (2013). Instant transformation of learned repulsion into motivational “wanting.” Current Biology, 23(4), 282–289. https://doi.org/10.1016/j.cub.2013.01.016

Robinson, T. E., & Flagel, S. B. (2009). Dissociating the predictive and incentive motivational properties of reward-related cues through the study of individual differences. Biological Psychiatry, 65(10), 869–873. https://doi.org/10.1016/j.biopsych.2008.09.006

Silva, F. J., Silva, K. M., & Pear, J. J. (1992). Sign-versus goal-tracking: Effects of conditioned-stimulus-to-unconditioned-stimulus distance. Journal of the Experimental Analysis of Behavior, 57(1), 17–31. https://doi.org/10.1901/jeab.1992.57-17

Skinner, B. F. (1948). Superstition in the pigeon. Journal of Experimental Psychology, 38(2), 168–172. https://doi.org/10.1037/h0055873

Starcke, K., Antons, S., Trotzke, P., & Brand, M. (2018). Cue-reactivity in behavioral addictions: A meta-analysis and methodological considerations. Journal of Behavioral Addictions, 7(2), 227–238. https://doi.org/10.1556/2006.7.2018.39

Villaruel, F. R., & Chaudhri, N. (2016). Individual differences in the attribution of incentive salience to a Pavlovian alcohol cue. Frontiers in Behavioral Neuroscience, 10. https://doi.org/10.3389/fnbeh.2016.00238

Williams, D. R., & Williams, H. (1969). Auto-maintenance in the pigeon: Sustained pecking despite contingent non-reinforcement. Journal of the Experimental Analysis of Behavior, 12(4), 511–520. https://doi.org/10.1901/jeab.1969.12-511

